# Spatial interactions between parrotfishes and implications for species coexistence

**DOI:** 10.1101/2022.10.21.513248

**Authors:** J.C. Manning, S.J. McCoy, S. Benhamou

## Abstract

Home range behavior mediates species interactions and distributions, and spatiotemporal segregation may facilitate coexistence of competing species. We investigated home range behavior and spatial interactions in four common parrotfishes on coral reefs in Bonaire, Caribbean Netherlands, to determine how spatial interactions mediate species interactions and contribute to their coexistence. We first computed home ranges for males and females of each species. We then quantified spatial overlap (i.e., static interaction) between the home ranges of neighboring male parrotfishes and their activity in shared areas to estimate interaction potential for pairs of individuals. Finally, we analyzed dynamic interactions in simultaneously tracked, spatially co-occurring interspecific pairs of parrotfishes to investigate how they interact in shared space. Generally, spatial overlap of home ranges was much lower for intraspecific pairs than for interspecific pairs, but the probability of finding males in areas shared with males of other species was species-dependent. Males in interspecific pairs moved mostly independently of each other in shared areas, but we did find some evidence of avoidance in interspecific pairs sharing the most space. We discuss our findings within the context of parrotfish social and foraging ecology to further elucidate the spatial ecology of these functionally important reef fishes.

## Introduction

In many species, individuals establish home ranges, broadly defined as self-restricted areas in which animals conduct their normal daily activities. These home ranges arise from individual movement decisions made by animals in response to their environment [1–3]. Home range characteristics (e.g., size) are determined by resource abundances [4], the space use of competitors [5,6], and intrinsic characteristics of the animal (e.g., locomotion and trophic status; Harestad & Bunnell 1979; Nash *et al.* 2015; Tamburello *et al.* 2015). Additionally, some animals defend all or a portion of their home ranges as territories, when the benefit of doing so outweighs the cost (Brown 1964; Kaufmann 1983).

Home range and territorial behaviors can affect population dynamics and regulation, including by setting upper limits on population size [12–15]. Furthermore, animal movement and space use can facilitate species coexistence (reviewed in [16]). Coexistence is maintained through mechanisms that minimize average fitness differences between organisms (equalizing) or increase the strength of intraspecific competition relative to interspecific competition (stabilizing; [17]). Spatial and temporal segregation, mediated by animal movement and activity patterns, may act as a stabilizing mechanism that facilitates coexistence of competing species [16,18–21]. Yet, these mechanisms remain largely unexplored in reef fish communities.

Parrotfishes act as important agents of bioerosion and sediment production, reworking, and transport on coral reefs [22–25]; and their grazing helps to maintain reef substrates in cropped, early successional states dominated by filamentous cyanobacteria and other microscopic photoautotrophs [26,27]. Epilithic and endolithic microautotrophs, including filamentous and mat-forming benthic cyanobacteria, are the primary nutritional targets of parrotfish foraging [26,28–30]. There is strong evidence that parrotfishes partition these resources at fine spatial scales by targeting substrata at different successional stages [28,31], and at broader scales by foraging in different habitat types (e.g., rubble v. high-relief habitat; [32]). In contrast, we know little about how movement and space use contribute to niche partitioning and coexistence in parrotfishes.

Adult male (terminal phase; TP) parrotfishes of many species defend stable, exclusive territories containing harems of intraspecific females (initial phase; IP) against intraspecific TPs, increasing their spawning success and access to high quality foods [33–36]. Territoriality may contribute to population regulation [12], and could impose spatial constraints on parrotfish foraging (i.e., focused grazing within core areas; [37]) that mediate spatial patterns of benthic community assembly [38]. Movement and space use may also influence competition and coexistence among parrotfishes. However, studies of parrotfish movement and space-use are often limited to correlating home range size to body size or resource abundance (e.g., [39,40]).

In this study, we investigated spatial interactions among four common Caribbean parrotfishes. Specifically, we tested the hypothesis that parrotfishes would avoid one another spatially and/or temporally to reduce competition for shared resources. To test this hypothesis, we estimated home ranges for several TP and IP fishes of each species to quantify differences in space use and the spatial overlap (i.e., static interaction) of co-occurring TP pairs (inter- and intraspecific). Furthermore, we analyzed the movements of simultaneously tracked interspecific pairs of TP parrotfishes to investigate the potential for dynamic interactions between these fishes in shared space. Finally, we conducted behavioral observations of agonistic interactions to provide context for our findings.

## Material and methods

### Study sites and data collection

We conducted our study in June-July 2021 at two fringing coral reef sites on the leeward coast of Bonaire, Caribbean Netherlands: Aquarius and Invisibles. These sites are characterized by relatively high coral cover and low macroalgal cover [30]. The abundance and biomass of different fish groups, including parrotfishes, are higher on Bonaire’s coral reefs relative to more heavily fished reefs in the Eastern Caribbean [41]. This is likely due to fisheries management efforts, including spear gun bans in 1971, the establishment of no fishing areas in 2008, bans on parrotfishes catches in 2010, and fish trap phase-outs in 2010 [42].

At each site, we collected two separate datasets: (1) GPS tracking and behavioral observations of several individuals, one by one, to estimate home ranges, quantify static interactions and activity in shared space, and explore the role of agonistic behavior in driving these patterns; and (2) simultaneous GPS tracking of pairs of neighboring parrotfishes to investigate dynamic interactions between spatially co-occurring individuals. GPS tracks were recorded using a Garmin GPSMAP 78sc by a snorkeler remaining at the surface, above the tracked individual, while a SCUBA diver monitored its behavior below the surface (except during dynamic interaction tracks; see below). Divers acclimated fish to their presence for ∼ 2 mins and subsequently observed fish behavior from ∼ 2 m to avoid influencing their behavior. Marine fishes in protected areas and areas with high diver visitation have reduced flight initiation distances [43]. Parrotfishes in Bonaire are, thus, likely to be habituated to divers because of long-standing spearfishing bans [42] and a thriving dive tourism industry.

### Space use, static interaction, and activity in shared space

At each study site, we tracked several TPs and IPs of four common parrotfishes, *Scarus taeniopterus*, *Scarus vetula*, *Sparisoma aurforenatum*, and *Sparisoma viride,* for ∼20 mins (Supplemental Information: Table S1). We attempted to track all territorial TPs within predetermined ∼1,000 m^2^ plots at each study site. We also tracked several haremic IP fishes from different territories for all species. We verified that our tracking duration was sufficient to capture diurnal (i.e., daytime) home range behavior by conducting visual assessments of stationarity (Figure S1; [44]). Individual parrotfishes are known to use these diurnal home ranges for extended periods of time [33,35,36].

For each fish, we computed home range and core areas, defined as the areas within the 95% and 50% cumulative isopleths of the utilization distribution, respectively, using movement-based kernel density estimation [45]. We then quantified the spatial overlap of home ranges (i.e., static interactions) for intra- and interspecific pairs of neighboring TPs using Bhattacharyya’s Affinity (BA; [46]). We also quantified the probability of finding each TP within the home range and core areas of neighboring fishes (i.e., the volume of the fish’s utilization distribution that lies within the home ranges and core areas of neighbors; [47]). Agonistic interactions between intra- and interspecific TPs occurred 2.66 ± 0.09 m (mean ± SE, n = 168) from the boundaries of TP home ranges (see below). We, therefore, defined neighboring TPs as individuals whose home range boundaries were within 2.66 m of one another. All utilization distributions, spatial overlaps, and probabilities of activity within home range and core areas were computed in the R package adehabitatHR [48].

All statistical models were fit in the R package glmmTMB [49]. We fit Gamma distributed generalized linear models with log-link functions to investigate differences in home range and core area size as a function of site, species, phase (TP or IP), log_10_ body mass, and the interaction of species and phase (Table S2). To assess differences in (1) spatial overlap of home ranges for pairs of neighboring TPs and (2) the probability of finding each TP in the home range and core areas of neighboring TPs, we fit zero-inflated beta regression models (Tables S3-S5). For each beta regression, we fit a full model including site, the number of days between the GPS tracks for each pair (fish were not all tracked the same day), and species pairing (e.g., *Sp. viride* – *Sc. vetula*) as fixed factors and a reduced model including only species pairing as a fixed factor. For the two models assessing the probability of finding TPs in the home ranges and core areas of neighboring TPs, we also included focal fish identification as a random effect. We conducted model selection (AICc) to determine whether to use the full or reduced model for each response variable. After determining what fixed effects to include in our models, we fit variable dispersion models (i.e., precision parameter allowed to vary as a function of explanatory variables, respectively) and used model selection (AICc) to determine whether a fixed or variable dispersion model fit best (i.e., a two-step selection process similar to [50]). The significance of the fixed effects in the conditional portion of the three final models was tested using Type III Wald’s χ^2^. We also computed marginal means for the conditional, zero-inflated, and dispersion components of each model using the R package emmeans [51].

### Spatial patterns of agonistic behavior

We concurrently video-recorded the behavior of each TP with a GoPro Hero 4 Silver (GoPro, Inc) and analyzed these videos in the behavioral analysis software BORIS [52]. We recorded the duration (to the closest second; those lasting less than 1 s were scored as 1 s) of all agonistic interactions (agonisms) and species and ontogenetic phase of the interactor. Apparent agonisms for which we were unable to identify the interactor were excluded from analyses. Video times were synchronized with GPS time to estimate where each agonism occurred. We assessed the predictors of agonism frequency using a generalized linear model fit to a negative binomial distribution (Table S6). We included site, focal species, and interactor identity (e.g., TP – intraspecific, etc.) as fixed effects and the log of observation time as an offset. We assessed the predictors of agonism duration (log transformed to meet the model’s assumptions) and the minimum distance between the agonism and home range boundary of the focal fish with a linear mixed model including site, focal species, and interactor identity as fixed effects and focal fish identification as a random effect (Tables S7 and S8). We graphically assessed model residuals to confirm that they met assumptions.

### Simultaneous tracking and dynamic interactions

We investigated dynamic interactions (i.e., the tendency of two animals to move together, to avoid each other or to move independently) between simultaneously tracked pairs of spatially co-occurring TP *Sp. viride* and *Sc. vetula* (Video 1) at both study sites (Table S9). We chose these species as they were the largest and easiest to track simultaneously with only snorkelers, and because they tend to forage on similar substrates [53]. As such, we hypothesized that individuals of these species would be most likely to avoid one another to prevent competition within shared areas. GPS tracks for these fishes lasted 31.2 ± 2.0 min (mean ± SD, *n* = 18) and were resampled to achieve synchronized relocations every 5s.

For each pair (n = 9), we identified the area of overlap between the two home ranges using the ‘st_intersection’ function in the R package sf [54]. Call *N_s,s_* the number of relocation pairs for which both individuals were in the shared area, and *N_s,e_*, *N_e,s_*, and *N_e,e_* the number of relocation pairs for which at least one of two individuals was in its respective exclusive area. We then contrasted the actual frequency *N_s,s_*/*T* with which both individuals of a given pair were in the shared area, where *T* = *N_s,s_*+*N_s,e_*+*N_e,s_*+*N_e,e_* is the total number of simultaneous relocation pairs, with the theoretical probability (*N_s,s_*+*N_s,e_*)(*N_s,s_*+*N_e,s_*)/*T^2^*that both individuals would simultaneously be in the shared area if they moved independently of each other. We then tested for avoidance or attractance within shared areas by comparing the difference in the observed and theoretical values with a Wilcoxon signed-ranks test. For each pair, we also quantified the spatial overlap of their home ranges and the probability of finding each fish in the home range and core area of the other (as described above). Finally, we investigated the tendency for each pair to move jointly, independently, or avoid one another when simultaneously present within the shared area by computing the dynamic interaction index [55] using the ‘IAB’ function in the R package wildlifeDI [56].

## Results

### Space use and static interactions

We found a significant interactive effect of species and phase on home range size (Wald’s χ^2^ = 14.01, df = 3, *p* = 0.003). There was only a marginal interactive effect of species and phase on the sizes of core areas (Wald’s χ^2^ = 6.86, df = 3, *p* = 0.076), but significant differences among species (Wald’s χ^2^ = 38.78, df = 3, *p* < 0.001). *Scarus taeniopterus* had the smallest home ranges and core areas of all species, while *Sc. vetula* and *Sp. viride* had the largest (Figure 1). Additionally, TP home ranges and core areas were larger than IP home ranges and core areas for all species except *Sc. taeniopterus* (Figure 1).

**Figure 1:**
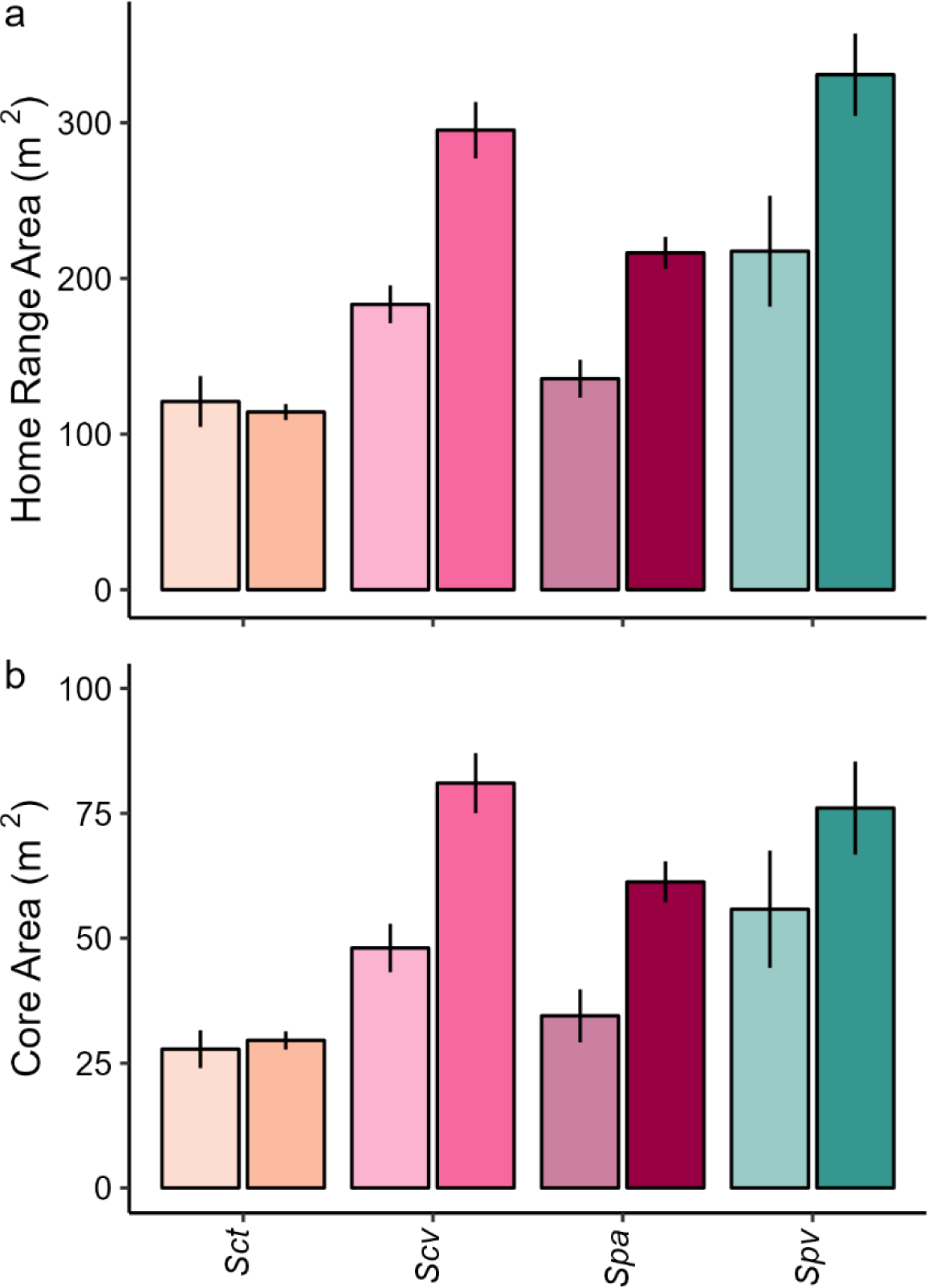
Mean (±SE) home range and core areas for TP (dark) and IP (light) of *Sc. taeniopterus*, *Sc. vetula*, *Sp. aurofrenatum*, and *Sp. viride*.

The best fit model for spatial overlap of neighboring TP parrotfishes was a zero-inflated beta regression with variable dispersion that included species pairings as an effect in each component of the model (Table S3). We found no spatial overlap between the home ranges of 16.7 – 36.7% pairs of neighboring TPs (Figure 2a). The likelihood of neighboring TP parrotfishes sharing space did not significantly differ among species pairs, but the home ranges of neighboring intraspecific pairs of TPs tended to be non-overlapping more often than the home ranges of neighboring interspecific pairs (Figure S2). When neighboring TP home ranges did overlap, the degree of overlap differed among species pairings (Wald’s χ^2^ = 112.01, df = 9, *p* < 0.001). For *Sc. vetula*, *Sp. aurofrenatum*, and *Sp. viride*, the home ranges of intraspecific TP pairs overlapped significantly less than the home ranges of interspecific TP pairs (Figure 2a). Overlap of intraspecific pairs of TP home ranges in *Sc. vetula*, *Sp. aurofrenatum*, and *Sp. viride* was extremely low (0.03 ± 0.01, 0.07 ± 0.02, and 0.04 ± 0.01; mean ± SE; *n* = 14, 25, and 11 pairs, respectively). For *Sc. taeniopterus*, there was no significant difference between the amount of space shared by intraspecific TP pairs and the amount of space shared with TP *Sp. viride* and TP *Sc. vetula* (Figure 2a and Figure S2).

**Figure 2:**
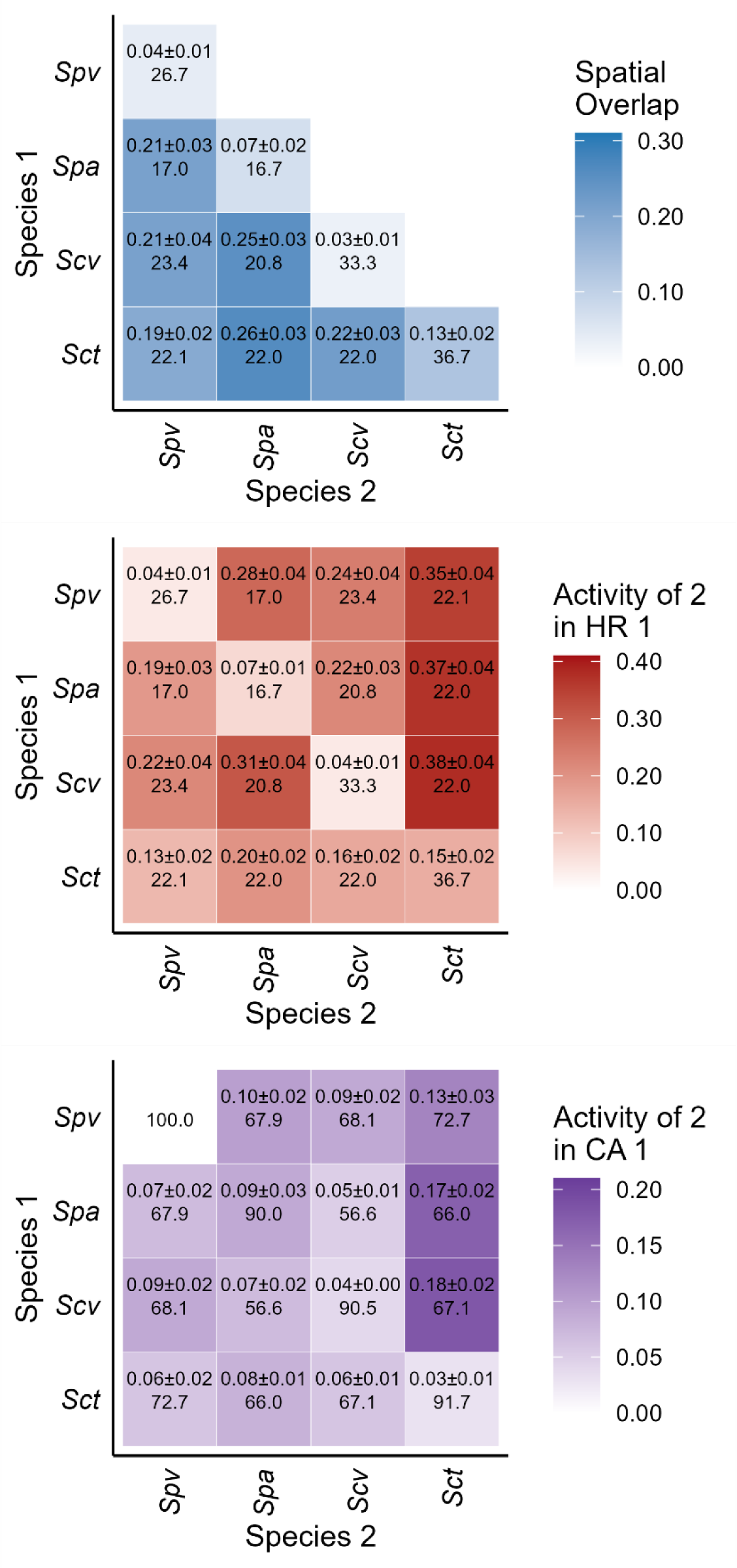
(a) A heatmap of the mean spatial overlap of the home ranges of neighboring TPs of *Sc. taeniopterus*, *Sc. vetula*, *Sp. aurofrenatum*, and *Sp. viride*. Lighter shades of blue indicate less spatial overlap. The mean (±SE) spatial overlap and the proportion of neighboring pairs with zero spatial overlap are displayed for each species pairing. (b) A heatmap of the mean probability of finding TPs of *Sc. taeniopterus*, *Sc. vetula*, *Sp. aurofrenatum*, and *Sp. viride* (Species 2) in the home ranges of neighboring TPs of *Sc. taeniopterus*, *Sc. vetula*, *Sp. aurofrenatum*, and *Sp. viride* (Species 1). Lighter shades of red indicate lower probabilities. The mean (±SE) probabilities and the proportions of neighboring pairs where the probability is zero are displayed for each species pairing. (c) A heatmap of the mean probability of finding TP of *Sc. taeniopterus*, *Sc. vetula*, *Sp. aurofrenatum*, and *Sp. viride* (Species 2) in the core areas of neighboring TPs of *Sc. taeniopterus*, *Sc. vetula*, *Sp. aurofrenatum*, and *Sp. viride* (Species 1). Lighter shades of purple indicate lower probabilities. The mean (±SE) probabilities and the proportions of neighboring pairs where the probability is zero are displayed for each species pairing.

The best fit model for the probability of finding focal TPs within the home ranges of neighboring TPs was a zero-inflated beta regression with variable dispersion that included species pairings as an effect in each component of the model and focal identity as a random effect (Table S4). The proportion of neighboring TP parrotfishes that shared no space (Figure 2b) did not differ among species pairings (Figure S3). When the home ranges of neighbors overlapped, the probability of finding focal TPs in the home ranges of their neighbors significantly differed among species pairings (Wald’s χ^2^ = 296.44, df = 15, *p* < 0.001). TP *Sc. vetula*, *Sp. aurofrenatum*, and *Sp. viride* were significantly less likely to be found in intraspecific home ranges than in interspecific home ranges (Figure 2b). The same was not true for TP *Sc. taeniopterus*. TP *Sc. taeniopterus* were more likely to be found in the home ranges of TP *Sc. vetula*, *Sp. aurofrenatum*, and *Sp. viride* than vice versa (Figure 2b and Figure S3).

The best fit model for the probability of finding focal TPs within the core areas of neighboring TPs was a zero-inflated beta regression with variable dispersion that included species pairings as an effect in each component of the model and focal identity as a random effect (Table S5). The proportion of focal TPs with zero probability of being found in the core areas of neighboring TPs was higher for intraspecific than interspecific pairs (Figure S4). In fact, the probability of finding neighboring TP *Sp. viride* in one another’s core areas was always zero (Figure 2c). When the probability of finding neighboring TPs in one another’s core areas was greater than zero, it was still low and differed significantly among species pairings (Wald’s χ^2^ = 231.68, df = 14, *p* < 0.001). TP *Sc. taeniopterus* had a higher probability of being found in the core areas of TP *Sc. vetula* and *Sp. aurofrenatum*, than the reverse (Figure 2c and Figure S4). Additionally, though not significantly, TP *Sc. taeniopterus* were also more likely to be found in the core areas of TP *Sp. viride* than the reverse (Figure 2c and Figure S4).

### Spatial patterns of agonisms

Focal TPs were most frequently agonistic toward IPs and TPs belonging to the same species (Wald’s χ^2^ = 160.76, df = 3, *p* < 0.001; Figure 3a). Intraspecific agonisms between TPs lasted longer than all other agonisms (Wald’s χ^2^ = 260.37, df = 3, *p* < 0.001; Figure 3b) and more frequently involved aggressive chases, rather than brief charges or displays (Fisher’s Exact Test p < 0.001). Intraspecific agonisms between focal TPs and IPs occurred 3.03 ± 0.07 m (mean ± SE, n = 292) from home range boundaries, further from the home range boundary than agonisms with other fishes (Wald’s χ^2^ = 11.40, df = 3, *p* = 0.010; Figure 3c). Focal TP *Sc. taeniopterus* engaged in agonisms more frequently than TPs of other species (Wald’s χ^2^ = 27.65, df = 3, *p* < 0.001; Figure S5a), and their agonisms occurred closer to home range boundaries (χ^2^ = 8.23, df = 3, *p* = 0.041; Figure S5b). Additionally, nearly all recorded interspecific agonisms (91.7%, n = 48) included TP *Sc. taeniopterus*, and the majority (52.3%) of these agonisms involved interactions between focal TP *Sc. taeniopterus* and IP *Sc. vetula*. TP *Sc. taeniopterus* were also involved in the majority (77.8%) of interspecific agonisms observed between TPs (n = 18). Of the interspecific agonisms recorded between TP *Sc. taeniopterus* and other TPs (n = 14), 50% were with TP *Scarus iseri*, 21.4% were with TP *Sc. vetula*, and the remaining were with TP *Sp. aurofrenatum* or TP *Sp. viride* (14.3% each).

**Figure 3:**
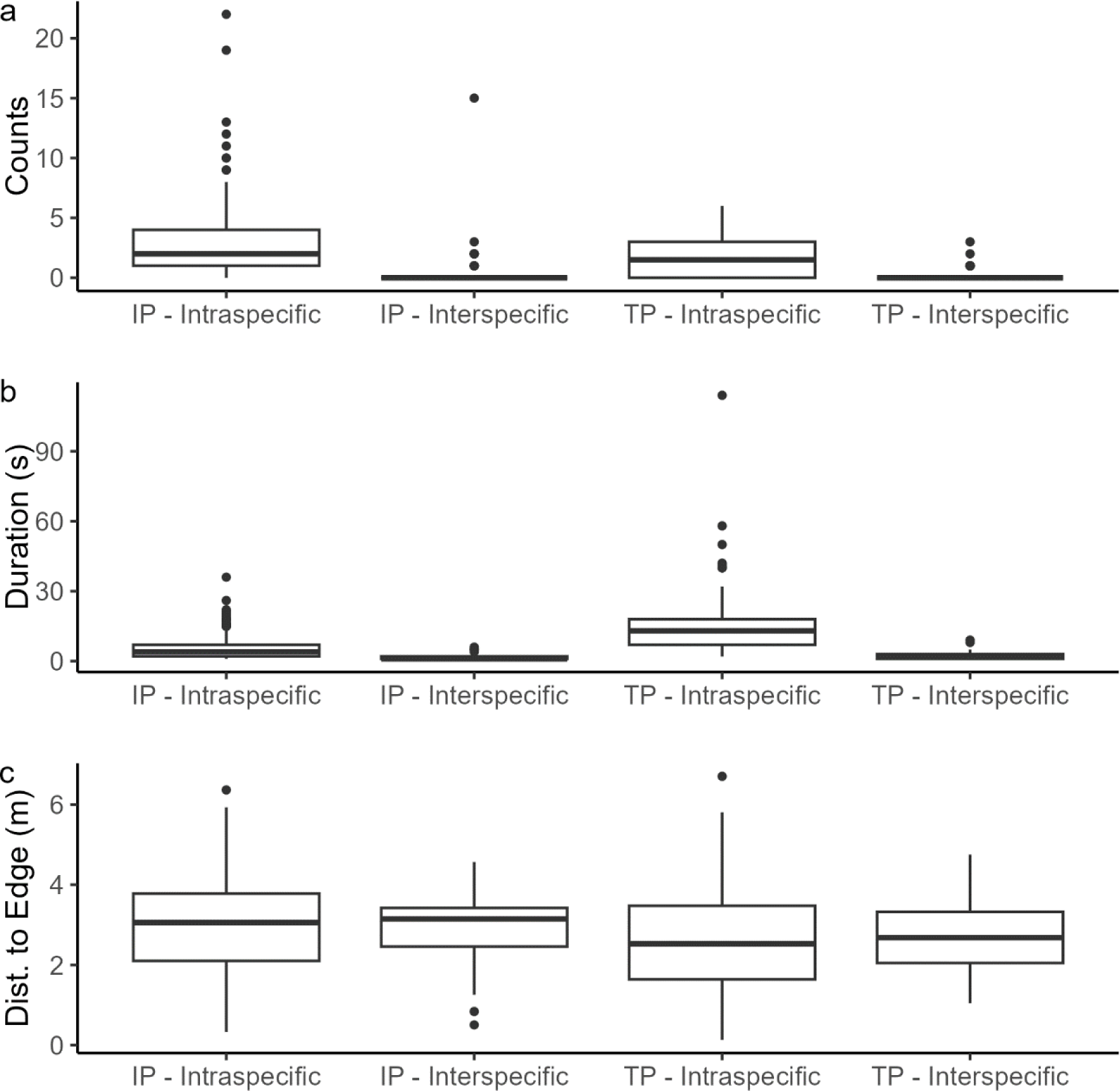
Boxplots of (a) the number of agonisms, (b) agonism durations (s), and (c) the distances (m) of those agonisms from the focal home range boundary for agonisms between focal TPs and IPs belonging to either the same species (IP-intraspecific) or another species (IP-interspecific), and TPs belonging either to the same species (TP-intraspecific) or another species (TP-interspecific).

### Dynamic interactions

The home ranges of simultaneously tracked interspecific pairs of TP *Sp. viride* and TP *Sc. vetula* overlapped substantially (0.51 ± 0.05, mean ± SE), providing a strong basis for potential dynamic interactions. However, the frequency of simultaneous relocations of these pairs within shared areas did not differ from the frequency expected for independent movement (*V* = 24, *p* = 0.91). We found statistically significant evidence of dynamic interaction in one pair of TP *Sp. viride* and TP *Sc. vetula* (Table S10; Video 1). This pair shared the most space (BA = 0.74) and exhibited significant avoidance while moving in shared areas (Table S10). Both individuals in this pair also had higher probabilities of being found in one another’s home ranges and core areas than individuals of most other pairs (Table S10). We also found marginally significant evidence of avoidance in another pair of TP *Sp. viride* and TP *Sc. vetula*, which had similarly high spatial overlap between their home ranges (BA = 0.64) and high probabilities of being found within one another’s home ranges and core areas (Table S10).

## Discussion

We quantified spatial interactions between TPs (i.e., males) of four parrotfish species in Bonaire to investigate the potential for movement and space use to mediate species interactions and coexistence in these functionally important reef fishes. Males of multiple parrotfish species are known to defend their daytime home ranges as fixed intraspecific territories from other males [33–35]. Consistent with prior work on parrotfish territoriality, we found strong intraspecific spatial segregation between neighboring TPs in *Sc. vetula, Sp. aurofrenatum,* and *Sp. viride,* likely driven by the aggressive territorial interactions we observed. Agonistic interactions generally occurred within 2-3 m of home range boundaries (Figure S6), suggesting that TPs quickly detected intruders within their territories. TP core areas lie almost entirely within these buffer zones (Figure S7), and the probability of finding neighboring TPs within core areas was zero for the majority of intraspecific pairs for these three species. Thus, TPs likely have near-exclusive access to the resources in these areas, aside from what they share with their harems of IP females.

Territorial behavior evolves in response to competition for limited resources, and is only tractable when resources are economically defendable (i.e., benefits of resource defense outweigh the costs; [10]). The density of competitors and resource availability, among other things, can strongly affect the cost-benefit balance of territory defense. Territory size is negatively related to competitor density in some species and experiments have demonstrated that removals of competitors may lead to territory expansion by remaining territory holders [34,57–59]. Competitor density also affects patterns of spatial overlap among territory holders [57,60], and direct encounters between intraspecific competitors can lead to home range shifts that persist for long periods of time [61]. *Scarus taeniopterus* is the most abundant parrotfish at our study sites [30]. The small home range sizes and increased intraspecific home range overlap that we found for TP *Sc. taeniopterus* relative to our other species could reflect trade-offs between territoriality and the density of intraspecific competitors. However, home range size also scales positively with body size in many animal taxa, including parrotfishes [7,9,40,62], and *Sc. taeniopterus* was the smallest of our study species. We cannot, therefore, rule out the effect of body size on these patterns, nor are we able to determine the effects of resource availability on observed spatial dynamics. Additional research is required to partition the effects of competitor density, resource dynamics, and allometry on spacing patterns in *Sc. taeniopterus* and other parrotfishes.

Resource partitioning is an important mechanism facilitating species coexistence. To date, much of the literature regarding resource partitioning in parrotfishes has centered on trophic resource partitioning. Microscopic photoautotrophs, including cyanobacteria, are the primary nutritional resources for parrotfishes [26]. Parrotfishes partition these resources at fine spatial scales by targeting substrates in different successional stages and with different species compositions [28,31,63]. *Sparisoma viride* and *Sc. vetula* primarily target sparse epilithic turfs growing atop endolithic communities, the former having increased access to endolithic communities as an excavating species [53,64,65]. Both species primarily target turfs on high-relief dead coral substrates, while *Sc. taeniopterus* primarily target turfs on boulder and rubble [32]. In contrast, *Sparisoma* spp., including *Sp. aurofrenatum* and *Sp. viride* regularly target fleshy macroalgae, though the macroalgae itself may not be the nutritional target [30,32,66]. While it is clear that parrotfishes partition trophic resources at various scales, it is less clear how spatial partitioning influences coexistence.

Spatial segregation can mitigate interference competition and act as a stabilizing mechanism of species coexistence [16]. For example, Berger and Gese [18] found that differential use of shared space, as evidenced by spatial interaction analyses, could promote coexistence in coyotes and wolves. Similarly, broad scale spatial segregation and fine scale patterns of avoidance, which vary seasonally, have been reported as potential mechanisms of coexistence in African carnivores and herbivores [20,67]. In this study, we found that the home ranges of neighboring interspecific pairs of TPs overlapped more than the home ranges of neighboring intraspecific pairs of TPs. However, spatial overlap was still relatively low (mean < 0.26), and the probability of finding neighboring TPs in one another’s home range and core areas varied by species. Specifically, TP *Sc. taeniopterus* were more likely be active within the home ranges and core areas of other species than vice versa, and were most active in the home ranges and core areas of TP *Sc. vetula*. TP *Sc. taeniopterus* were also involved in the majority of the interspecific agonistic interactions that we observed, mostly interacting with IP *Sc. vetula*. Haremic IPs share a substantial amount of space with and are very active in the TP home ranges and core areas to which they belong (Section S1, Table S11, Figures S8 and S9). Frequent interactions between TP *Sc. taeniopterus* and IP *Sc. vetula* likely reflect competition for shared resources in the space they share with TP *Sc. vetula*. In contrast, TP *Sc. taeniopterus* were nearly as likely to be found in neighboring TP *Sp. aurofrenatum* and TP *Sp. viride* home range and core areas, but were less likely to interact agonistically with these species. This may reflect a greater degree of trophic resource partitioning between *Sc. taeniopterus* and these other species, and reduced importance of spatial segregation to their coexistence. These results suggest that spatial partitioning may limit agonistic interactions between species, particularly when trophic resource partitioning is not sufficient to prevent competition for shared resources.

Animals can also partition space temporally by avoiding each other when present in shared space, thereby limiting interference competition (e.g., [68]). We detected significant avoidance in a single interspecific pair of TP *Sp. viride* and TP *Sc. vetula*. The home ranges of this pair had the greatest degree of spatial overlap of all simultaneously tracked pairs of TP *Sp. viride* and TP *Sc. vetula*. Likewise, we detected marginal avoidance in another pair of TP *Sp. viride* and TP *Sc. vetula* that also shared a substantial amount of space, with the second highest measure of home range overlap. It seems reasonable to assume that neighboring individuals of different parrotfish species may interact dynamically and avoid one another in shared space dependent upon the degree of home range overlap, but that under typical conditions spatial overlap may be low enough to limit the need for active avoidance. It is also possible that dynamic avoidance takes place at smaller spatiotemporal scales. In the future, it may be necessary to investigate dynamic interactions more locally to elucidate the underlying drivers of dynamic interactions within shared space [56]. In light of our findings that TPs of *Sc. taeniopterus* were more likely to be found within the home ranges and core areas of interspecific neighbors than other species, investigations of dynamic interactions between TP *Sc. taeniopterus* and other TP parrotfishes may yield stronger evidence of temporal segregation within shared space for Caribbean parrotfishes.

Home range sizes are correlated with the abundance and quality of resources [69,70], including for parrotfishes [39], and animals are predicted to increase the size of their home ranges in less productive habitats [8]. Anthropogenic stressors, including climate change, are leading to rapid and dramatic shifts in the structure and composition of benthic communities on coral reefs [71]. As these changes continue, we might expect the potential for spatial interactions to change as home ranges expand, contract, and/or shift in response to changes in resource availability. Baseline studies of spatial interactions such as this are, therefore, increasingly important for understanding the effects of climate change on reef fish communities [72].

The results presented here elucidate a few of the mechanisms underlying movement and space use in parrotfishes. Specifically, we found that competition drove stronger spatial segregation between TPs belonging to the same species than to different species. Dynamic avoidance does occur between TPs belonging to different species, though such interactions may depend on how much space each member of a pair shares, and how such shared space is used. Regardless the presence of dynamic avoidance suggests that spatiotemporal segregation could act as a stabilizing mechanism mediating coexistence in parrotfishes. Our results have implications for understanding the trophic effects of parrotfishes on coral reefs. The effects of grazers on autotrophs depends, in part, on spatial variation in grazing pressure [73]. Home range and territorial behavior may concentrate parrotfish grazing and bioerosion locally [37], contributing to spatial heterogeneity in benthic reef communities. As such, studies of the spatial ecology of parrotfishes, particularly those that investigate spatial interactions, are necessary to provide a more complete understanding of their functional roles on coral reefs.

## Acknowledgements and funding statement

This material is based upon research supported by the Chateaubriand Fellowship of the Office for Science & Technology of the Embassy of France in the United States. The research was also supported by Florida State University (FSU) start-up funding and the Tatelbaum Ocean Research Fund awarded to SJM, as well as a Mote Research Assistantship and Jack Winn Gramling Research Award from FSU, a Guy Harvey Scholarship, and a Society for Integrative and Comparative Biology Grant-In-Aid of Research awarded to JCM. We would like to thank R. Francisca and C. Eckrich at Stinapa Bonaire for assistance with permitting to work in the Bonaire National Marine Park. We are also grateful for the valuable feedback that E. Cissell and A. Rassweiler provided on earlier versions of this manuscript. Finally, we thank L. Kury and K. Laforest for assisting with data collections.

