## Supplemental Information for "Spatial interactions between parrotfishes and implications for species coexistence"

319 Stadium Drive

Tallahassee, FL 32306-4295, USA

[2] University of North Carolina, Department of Biology

120 South Road, CB3280

Chapel Hill, NC 27599-3280, USA

[3] MAD Team, Centre d'Ecologie Fonctionnelle et Evolutive, CNRS

1919 Route de Mende, 34293 Montpellier Cedex 5, France

Associated to Cogitamus Lab

Table S1: Number of tracks (*N*) and mean ± SD of track duration and number of GPS relocations obtained per track for each species and ontogenetic phase.

| **Species** | **Phase** | ***N*** | **Duration (min)** | **Relocations** |
|---|---|---|---|---|
| *Sc. taeniopterus* | IP | 11 | 18.97 ± 5.32 | 113.36 ± 31.79 |
| | TP | 35 | 20.09 ± 2.05 | 115.49 ± 23.03 |
| *Sc. vetula* | IP | 11 | 20.38 ± 1.06 | 121.45 ± 24.87 |
| | TP | 17 | 22.47 ± 1.68 | 130.94 ± 14.79 |
| *Sp. aurofrenatum* | IP | 10 | 20.50 ± 0.46 | 119.20 ± 18.75 |
| | TP | 20 | 19.81 ± 3.86 | 122.05 ± 29.36 |
| *Sp. viride* | IP | 10 | 21.48 ± 2.92 | 118.70 ± 13.92 |
| | TP | 14 | 21.15 ± 0.96 | 120.57 ± 23.20 |

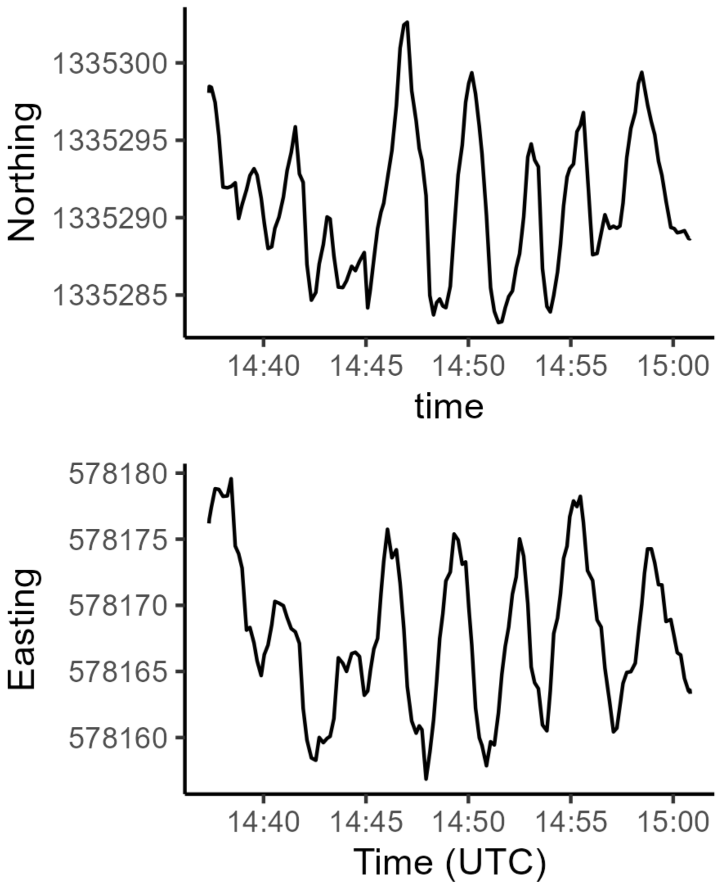


Figure S1: Example time series plot demonstrating the locational stationarity of a TP *Sc. vetula* at Invisibles: the mean and variance of location coordinates does not seem to vary across time (e.g., between the first and second halves of the time series)

Table S2: Output of generalized linear models (Gamma distribution) of home range and core areas ($m^2$) as a function of site, species, ontogenetic phase, $\log_{10}$(body mass), and the interaction between species and ontogenetic phase.

| | | | | |
|---|---|---|---|---|
| **Home range** | | | | |
| | **Wald's $\chi^2$** | **df** | ***p*-value** | |
| Intercept | 34.93 | 1 | < 0.001 | *** |
| Site | 2.11 | 1 | 0.147 | |
| Species | 53.33 | 3 | < 0.001 | *** |
| Phase | 1.45 | 1 | 0.228 | |
| $\log_{10}$(body mass) | 2.10 | 1 | 0.148 | |
| Species:Phase | 14.01 | 3 | 0.003 | ** |
| **Core area** | | | | |
| | **Wald's $\chi^2$** | **df** | ***p*-value** | |
| Intercept | 9.38 | 1 | 0.002 | ** |
| Site | 1.41 | 1 | 0.235 | |
| Species | 38.78 | 3 | < 0.001 | *** |
| Phase | 1.44 | 1 | 0.231 | |
| $\log_{10}$(body mass) | 1.00 | 1 | 0.318 | |
| Species:Phase | 6.86 | 3 | 0.076 | |

Table S3: Model selection and output of the conditional model for home range overlap estimated using Bhattacharyya's Affinity. Best fit model is indicated in bold italics.

| Model | Conditional | Zero-Inflation | Dispersion | Random | df | AICc |
|---|---|---|---|---|---|---|
| Full | Site + Interaction Type + Days b/w Tracks | Site + Interaction Type + Days b/w Tracks | | 1\|ID | 26 | 10.11 |
| Reduced | Interaction Type | Interaction Type | | 1\|ID | 22 | 5.82 |
| ***Var. Dispersion*** | ***Interaction Type*** | ***Interaction Type*** | ***Interaction Type*** | ***1\|ID*** | ***31*** | ***-19.58*** |

| | Wald's $\chi^2$ | df | *p*-value | |
|---|---|---|---|---|
| Intercept | 812.39 | 1 | < 0.001 | *** |
| Interaction Type | 112.01 | 9 | < 0.001 | *** |
| **Random Effects** | **Variance** | **SD** | | |
| Individual | $1.14 \times 10^{-10}$ | $1.07 \times 10^{-5}$ | | |

Table S4: Model selection and output of the conditional model for the probability of finding each TP in the home ranges of neighboring TPs. Best fit model is indicated in bold italics.

| Model | Conditional | Zero-Inflation | Dispersion | Random | df | AICc |
|---|---|---|---|---|---|---|
| Full | Site + Interaction Type + Days b/w Tracks | Site + Interaction Type + Days b/w Tracks | | 1\|ID | 38 | 147.43 |
| Reduced | Interaction Type | Interaction Type | | 1\|ID | 34 | 147.48 |
| ***Var. Dispersion*** | ***Interaction Type*** | ***Interaction Type*** | ***Interaction Type*** | ***1\|ID*** | ***49*** | ***19.27*** |

| | Wald's $\chi^2$ | df | *p*-value | |
|---|---|---|---|---|
| Intercept | 1179.50 | 1 | < 0.001 | *** |
| Interaction Type | 296.44 | 15 | < 0.001 | *** |
| | **Variance** | **SD** | | |
| Individual | $6.94 \times 10^{-10}$ | $2.63 \times 10^{-5}$ | | |

Table S5: Model selection and output of the conditional model for the probability of finding each TP in the core areas of neighboring TPs. Best fit model is indicated in bold italics.

| Model | Conditional | Zero-Inflation | Dispersion | Random | df | AICc |
|---|---|---|---|---|---|---|
| Full | Site + Interaction Type + Days b/w Tracks | Site + Interaction Type + Days b/w Tracks | | 1\|ID | 36 | 400.50 |
| Reduced | Interaction Type | Interaction Type | | 1\|ID | 32 | 395.18 |
| ***Var. Dispersion*** | ***Interaction Type*** | ***Interaction Type*** | ***Interaction Type*** | ***1\|ID*** | ***46*** | ***383.18*** |

| | Wald's $\chi^2$ | df | *p*-value | |
|---|---|---|---|---|
| Intercept | 2024.09 | 1 | < 0.001 | *** |
| Interaction Type | 231.68 | 14 | < 0.001 | *** |
| | **Variance** | **SD** | | |
| Individual | $1.54\times10^{-10}$ | $1.24\times10^{-5}$ | | |

Table S6: Output of a generalized linear model of agonism frequency fit to a negative binomial distribution, with site, species, and identity of the interactor as fixed effects and the log of observation time as an offset in the model.

| | **Wald's $\chi^2$** | **df** | ***p*-value** | |
|---|---|---|---|---|
| Intercept | 1080.78 | 1 | < 0.001 | *** |
| Site | 0.23 | 1 | 0.634 | |
| Species | 27.65 | 3 | < 0.001 | *** |
| Interactor ID | 160.76 | 3 | < 0.001 | *** |

Table S7: Output of the linear mixed model of $\log_{10}$ transformed agonism durations (s) as a function of site, species, and interactor identity, with focal individual identity as a random effect.

| | **Wald's $\chi^2$** | **df** | ***p*-value** | |
|---|---|---|---|---|
| Intercept | 338.17 | 1 | < 0.001 | *** |
| Site | 3.37 | 1 | 0.066 | |
| Species | 4.75 | 3 | 0.191 | |
| Interactor ID | 260.37 | 3 | < 0.001 | *** |
| | **Variance** | **SD** | | |
| Individual | 0.008 | 0.089 | | |
| Residual | 0.096 | 0.310 | | |

Table S8: Output of the linear mixed model of the distances of agonisms from the edge of the home range boundary (m) as a function of site, species, and interactor identity, with focal individual identity as a random effect.

| | **Wald's $\chi^2$** | **df** | ***p*-value** | |
|---|---|---|---|---|
| Intercept | 678.97 | 1 | < 0.001 | *** |
| Site | 0.47 | 1 | 0.494 | |
| Species | 8.23 | 3 | 0.041 | * |
| Interactor ID | 11.40 | 3 | 0.010 | ** |
| | **Variance** | **SD** | | |
| Individual | 0.080 | 0.283 | | |
| Residual | 1.288 | 1.135 | | |

Table S9: Number of tracks (*N*) and mean ± SD home range area, track duration, and number of relocations per track for simultaneously tracked TP *Sc. vetula* and TP *Sp. viride*.

| Focal Species | Focal Phase | N | Relocations | Duration (min) | Home range area ($m^2$) |
|---|---|---|---|---|---|
| *Sc. vetula* | TP | 9 | 183.78 ± 41.50 | 31.75 ± 2.44 | 307.17 ± 55.66 |
| *Sp. viride* | TP | 9 | 175.56 ± 22.11 | 30.69 ± 1.39 | 389.25 ± 87.16 |

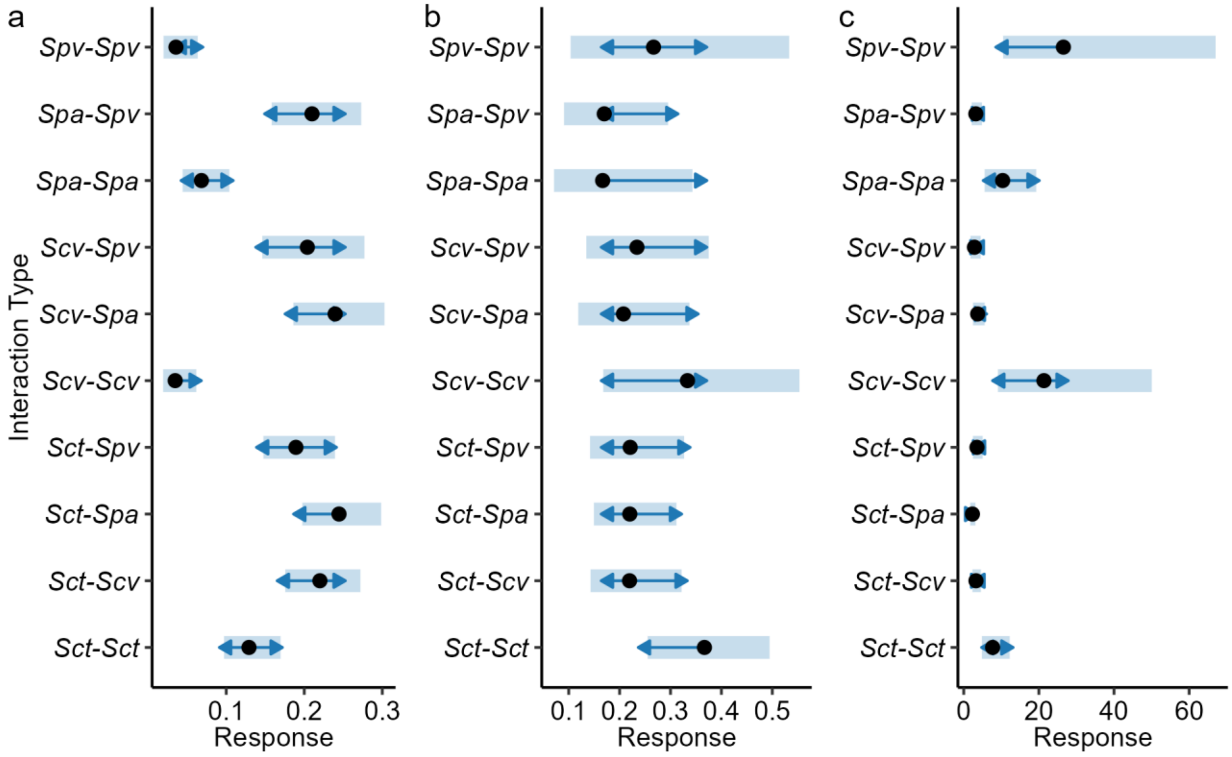


Figure S2: Estimated marginal means (± 95% asymptotic CIs) for the (a) conditional, (b) zero-inflated, and (c) dispersion components of the zero-inflated beta regression of spatial overlap among neighboring TP parrotfishes. Non-overlapping arrows represent significant differences among groups.

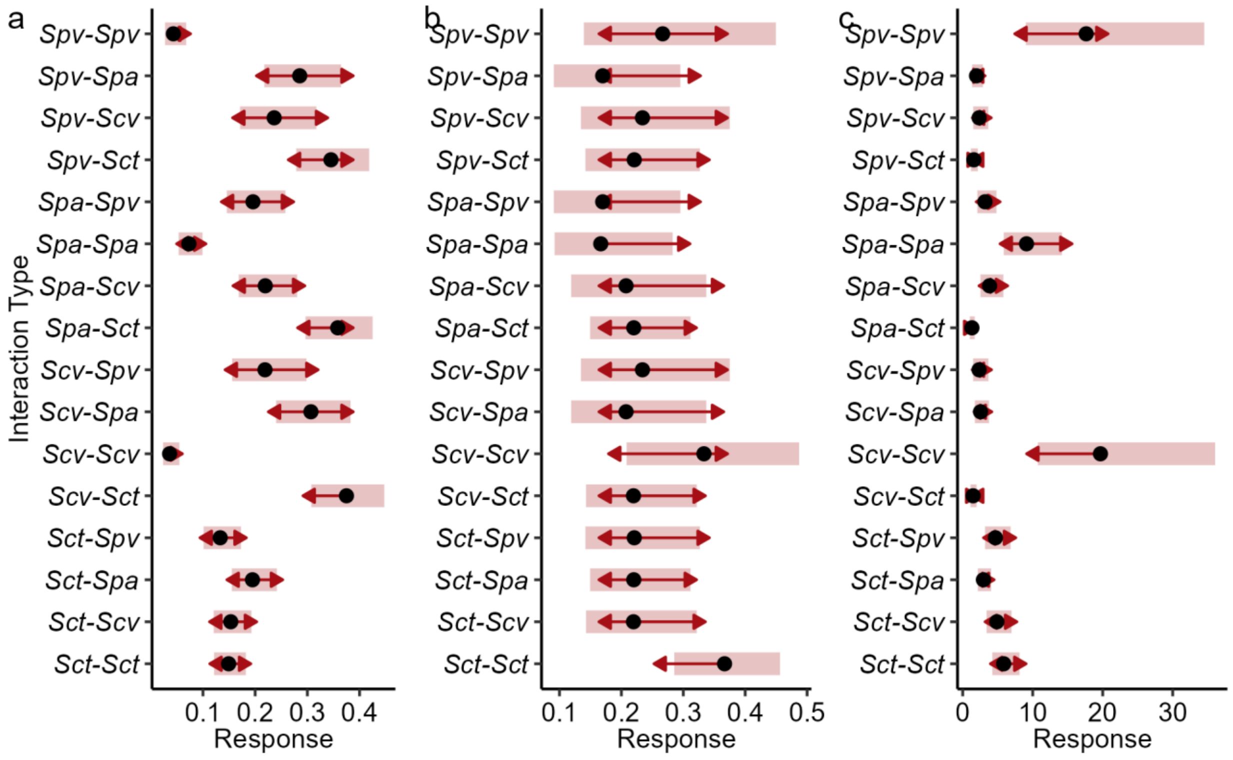


Figure S3: Estimated marginal means (± 95% asymptotic CIs) for the (a) conditional, (b) zero-inflated, and (c) dispersion components of the zero-inflated beta regression of activity within the home ranges of neighboring TP parrotfishes. Non-overlapping arrows represent significant differences among groups.

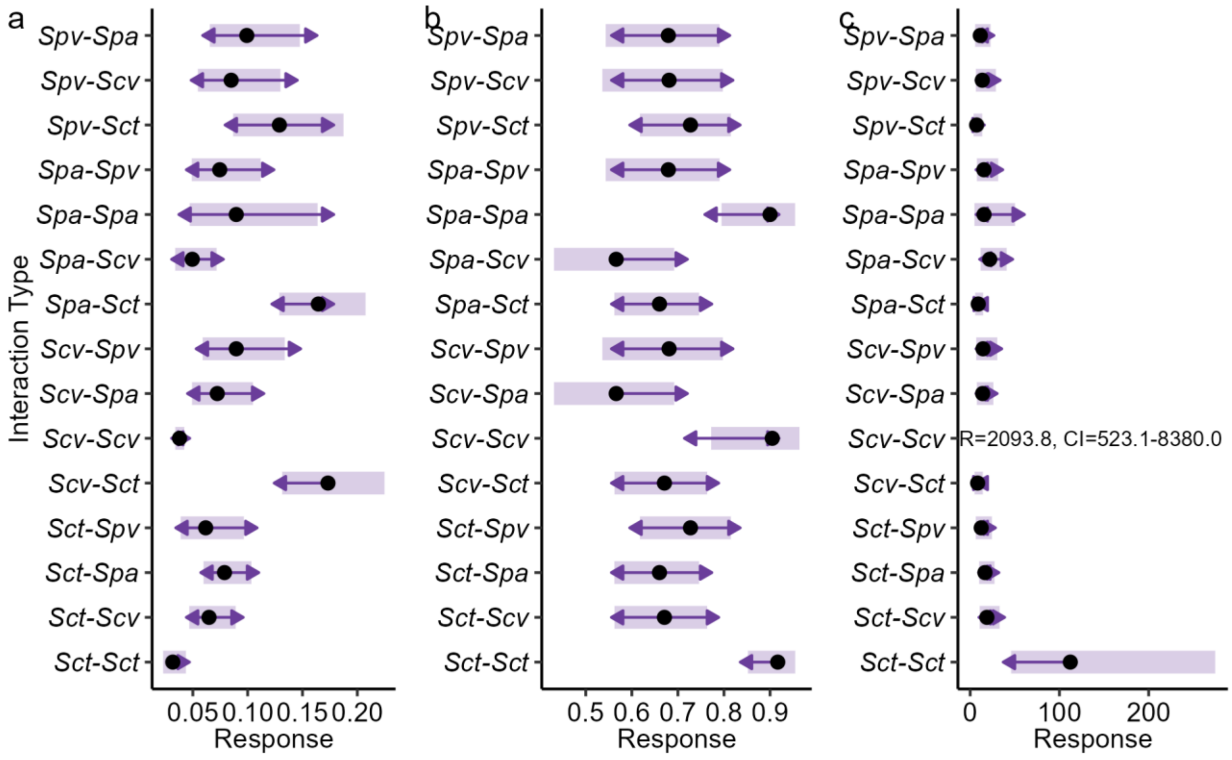


Figure S4: Estimated marginal means (± 95% asymptotic CIs) for the (a) conditional, (b) zero-inflated, and (c) dispersion components of the zero-inflated beta regression of activity within the core areas of neighboring TP parrotfishes. Non-overlapping arrows represent significant differences among groups. Per warning message when plotting estimated marginal means, arrows are not reflective of significance for comparisons between two pairs: Sct-Sct and Sct-Spv; Sct-Sct and Spv-Sct. Their arrows should not overlap but do (target overlap = $-1x10^{-4}$, overlap on graph = $5x10^{-4}$ for both).

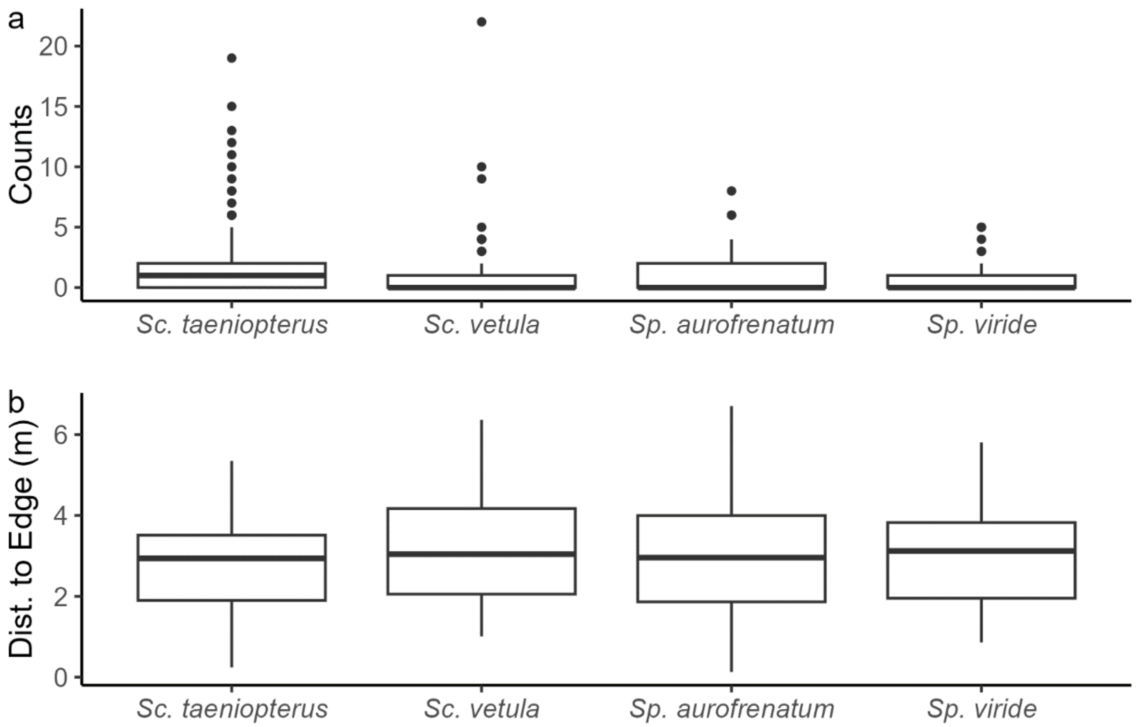


Figure S5: Boxplots of (a) the number of agonisms and (b) the distances of those agonisms from the boundary between focal terminal phase parrotfishes and other parrotfishes for *Sc. taeniopterus*, *Sc. vetula*, *Sp. aurofrenatum*, and *Sp. viride*.

Table S10: Overlap of home ranges (Bhattacharyya's Affinity), the probability for each individual to be found within the other individual's home range and core area, and observed and expected (under the null hypothesis of independent movements) dynamic interaction level, and significance values for one-tailed tests of attractance and avoidance for interspecific pairs of parrotfishes.

| **Type** | **Site** | **HR Overlap** | **Activity of 1 in 2's HR** | **Activity of 2 in 1's HR** | **Activity of 1 in 2's CA** | **Activity of 2 in 1's CA** | **Observed Dynamic Interaction** | **Expected Dynamic Interaction** | **Attract *p*-value** | **Avoid *p*-value** |
|---|---|---|---|---|---|---|---|---|---|---|
| Interspecific<br>1 TP *Sp. viride*<br>2 TP *Sc. vetula* | AQ | 0.273 | 0.318 | 0.266 | 0.048 | 0.061 | 0.871 | 0.892 | - | 0.385 |
| | | 0.467 | 0.334 | 0.653 | 0.075 | 0.120 | 0.828 | 0.844 | - | 0.357 |
| | | 0.552 | 0.563 | 0.639 | 0.059 | 0.064 | 0.683 | 0.639 | 0.277 | - |
| | | **0.741** | **0.692** | **0.875** | **0.171** | **0.315** | **0.613** | **0.681** | - | **0.004** |
| | | 0.466 | 0.363 | 0.624 | 0.085 | 0.116 | 0.734 | 0.728 | 0.383 | - |
| | IV | 0.398 | 0.354 | 0.488 | 0.087 | 0.144 | 0.753 | 0.778 | - | 0.244 |
| | | ***0.642*** | ***0.729*** | ***0.625*** | ***0.172*** | ***0.137*** | ***0.595*** | ***0.673*** | - | ***0.070*** |
| | | 0.555 | 0.521 | 0.579 | 0.190 | 0.199 | 0.834 | 0.788 | 0.108 | - |
| | | 0.487 | 0.400 | 0.684 | 0.025 | 0.025 | 0.711 | 0.747 | - | 0.161 |

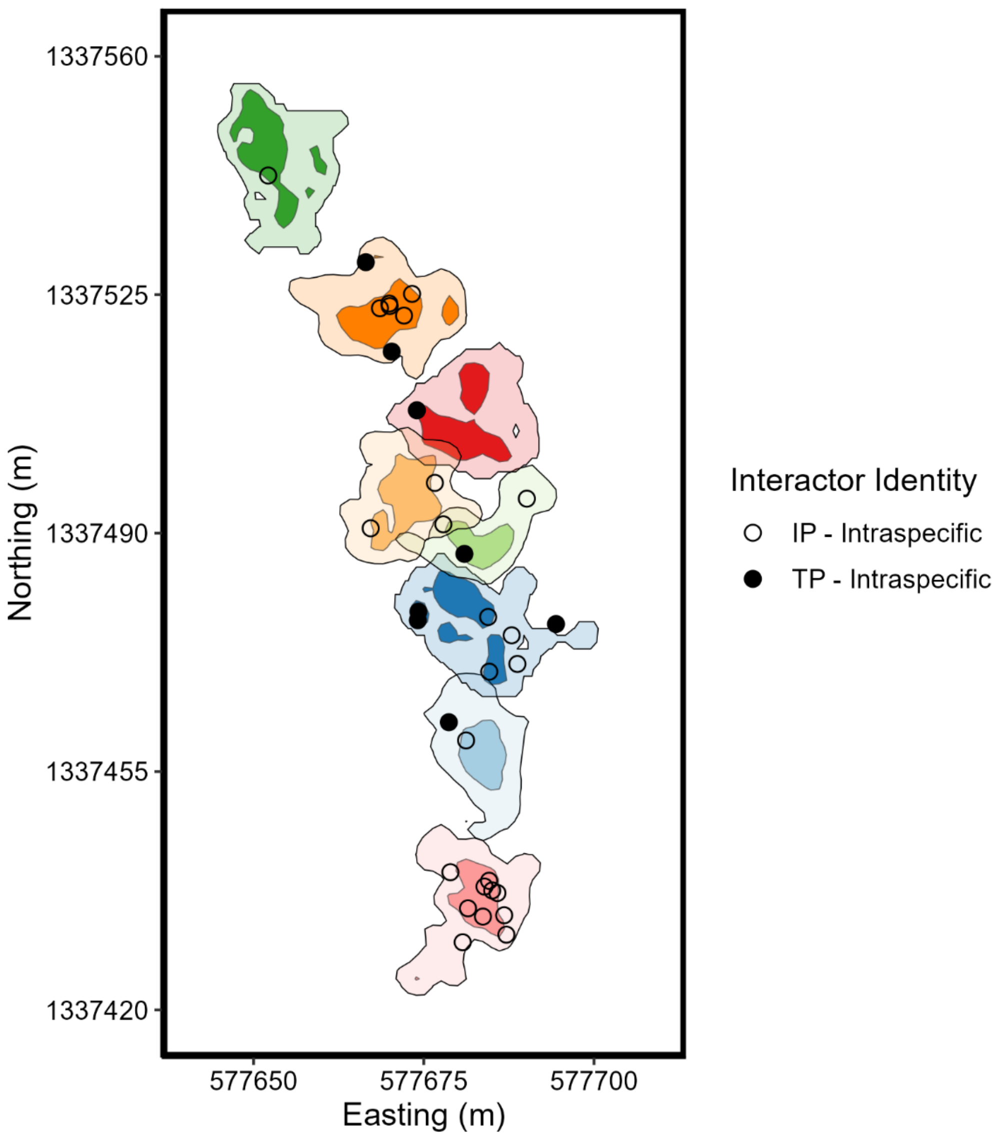


Figure S6: TP *Sc. vetula* home ranges and core areas (shaded darker) at Aquarius fringing reef, Bonaire. The home ranges of different individuals are shown in different colors. The points show the locations of intraspecific agonisms with IP (open circles) and TP (closed circles) individuals. Mapped using the UTM 19N (EPSG: 32619) projection.

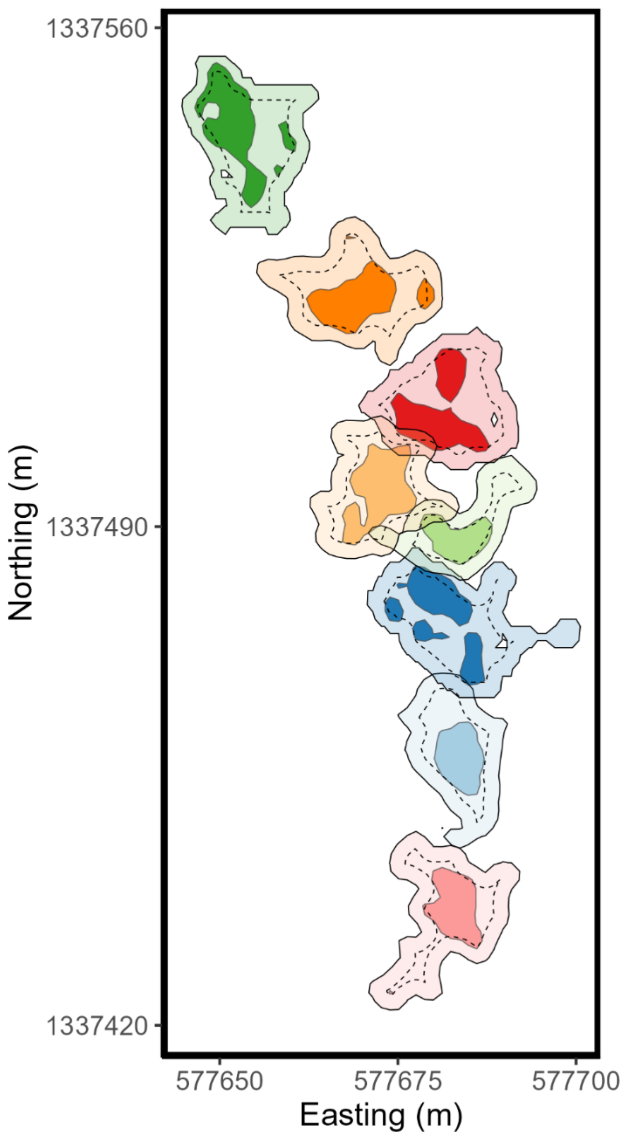


Figure S7: TP *Sc. vetula* home ranges and core areas (shaded darker) at Aquarius fringing reef, Bonaire. The home ranges of different individuals are shown in different colors. The dashed lines within each home range denote the distance from the home range boundary where intraspecific TP agonisms occurred on average (2.13 m for *Sc. vetula*). Mapped using the UTM 19N (EPSG: 32619) projection.

Section S1: Spatial interactions between haremic pairs of TP and IP parrotfish

We quantified the spatial overlap of TP and IP home ranges for each species using Bhattacharyya's Affinity (BA), and a given IP was assumed to belong to the TP's harem with which it shared the most space. The utilization distributions and spatial overlaps were computed in the adehabitatHR R package [1]. To assess differences in the spatial overlap of haremic TP and IP home ranges among species, we fit beta regression models. We fit a full model including site, the number of days between the GPS tracks for each pair (fish were not all tracked the same day), and species as fixed factors and a reduced model including only species as a fixed factor. We conducted model selection (AICc) to determine whether to use the full or reduced model for each response variable. After determining what fixed effects to include in our models, we fit variable dispersion models (i.e., precision parameter allowed to vary as a function of explanatory variables, respectively) and used model selection (AICc) to determine whether a fixed or variable dispersion model fit best (i.e., a two-step selection process similar to [2]). All statistical models were fit in the R package glmmTMB [3]. The significance of the fixed effects in the conditional portion of the three final models was tested using Type III Wald's $\chi^2$.

The best fit model for spatial overlap of home ranges for haremic TP and IP pairs was a beta regression with fixed dispersion that included species as a fixed factor in the model (Table S11). There were no significant differences among species in how much space was shared by haremic TP and IP pairs (Wald's $\chi^2 = 2.69$, df = 3, $p = 0.443$; Figure S8). Home range overlap was high for all species (0.52 ± 0.03, mean ± SE, n = 41 pairs).

Table S11: Output for conditional model of spatial overlap of home ranges for TP and IP pairs (Bhattacharyya's Affinity).

| **Model** | **Conditional** | **Dispersion** | **df** | **AICc** |
|---|---|---|---|---|
| Full | Site + Species + Days b/w Tracks | | 7 | -5.29 |
| ***Reduced*** | ***Species*** | | ***5*** | ***-7.43*** |
| Var. Dispersion | Species | Species | 8 | -1.36 |

| | **Wald's $\chi^2$** | **df** | ***p*-value** |
|---|---|---|---|
| Intercept | 0.34 | 1 | 0.558 |
| Species | 2.69 | 3 | 0.443 |

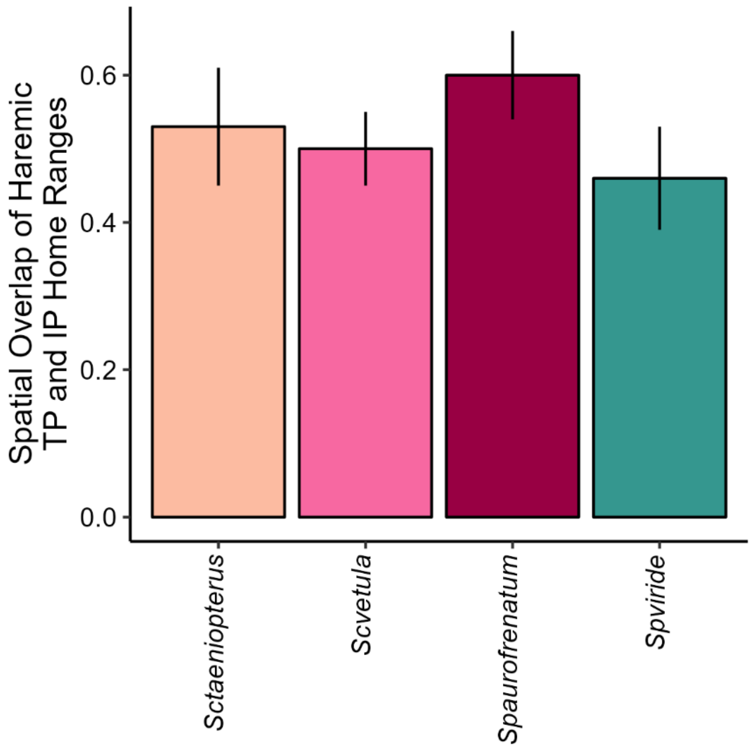


Figure S8: Mean (±SE) home range overlap (BA) for haremic TP and IP pairs of *Sc. taeniopterus*, *Sc. vetula*, *Sp. aurofrenatum*, and *Sp. viride*.

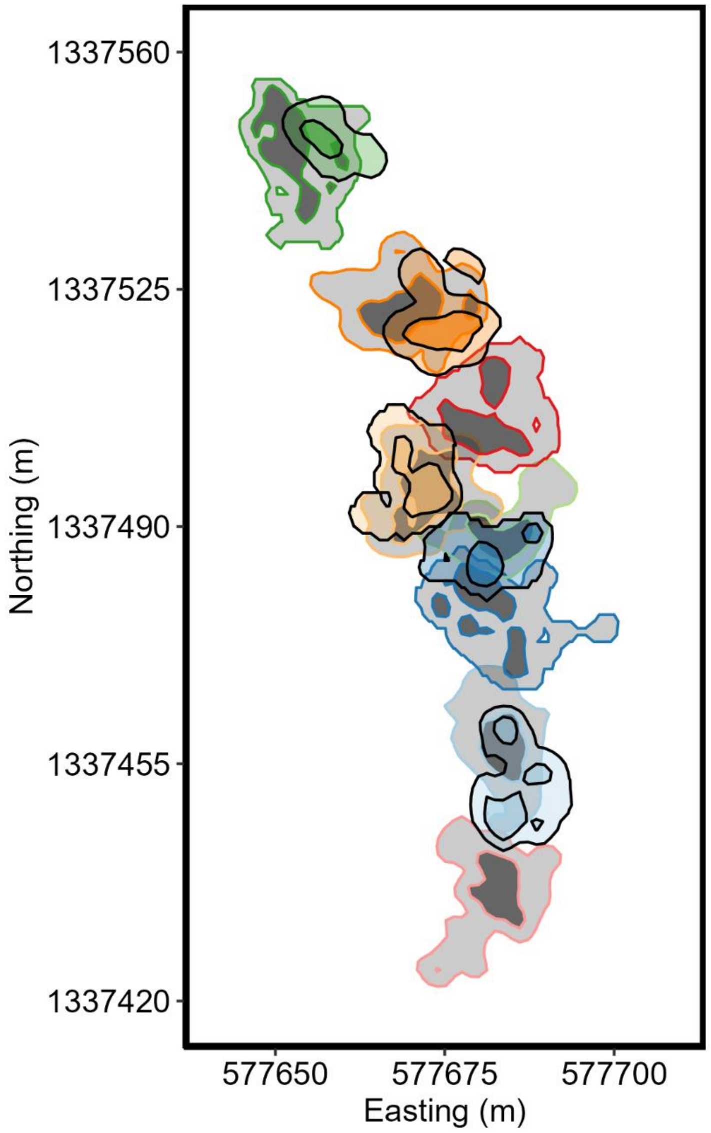


Figure S9: TP and IP *Sc. vetula* home ranges and core areas (shaded darker) at Aquarius fringing reef, Bonaire. TP home ranges and core areas are filled in gray scale and outlined in colors specific to each individual. IP home ranges and core areas are filled with the color that matches the color of the TP home range to which it was assigned. Mapped using the UTM 19N (EPSG: 32619) projection.
