## Supplementary material for "Spatial interactions between parrotfishes and implications for species coexistence": Video 1

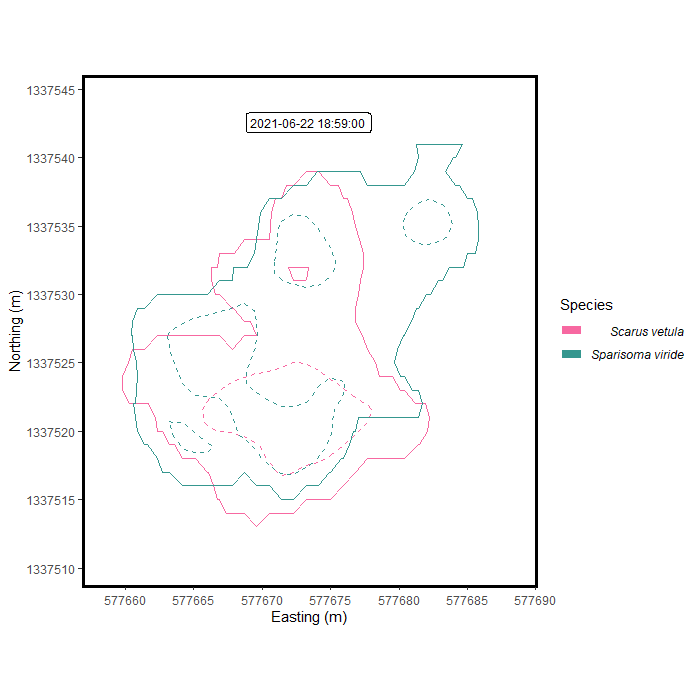

Video 1: Movements of a simultaneously tracked pair of TP *Sp. viride* (dark green) and TP *Sc. vetula* (pink) at Aquarius fringing reef, Bonaire, mapped using the UTM 19N (EPSG: 32619) projection. The home ranges (solid lines) and core areas (dashed lines) represent the 95% and 50% cumulative isopleths, respectively, of the utilization distributions computed for the TP *Sp. viride* (dark green) and TP *Sc. vetula* (pink) using movement-based kernel density estimation. The time is presented in UTC. This pair exhibited significant dynamic avoidance in shared areas of their home ranges.
